# Oncolytic adenovirus serotype 35 mediated tumor growth suppression *via* efficient activation and tumor infiltration of natural killer cells

**DOI:** 10.1101/2022.12.09.519732

**Authors:** Ryosuke Ono, Fuminori Sakurai, Ken J. Ishii, Hiroyuki Mizuguchi

**Author notes:** **Correspondence:** Fuminori Sakurai, Laboratory of Biochemistry and Molecular Biology, Graduate School of Pharmaceutical Sciences, Osaka University, 1-6 Yamadaoka, Suita-city, Osaka 565-0871, Japan., Hiroyuki Mizuguchi, Laboratory of Biochemistry and Molecular Biology, Graduate School of Pharmaceutical Sciences, Osaka University, 1-6 Yamadaoka, Suita-city, Osaka 565-0871, Japan.

## Abstract

**Background:** Oncolytic adenoviruses (OAds) mediate superior antitumor effects both by inducing direct oncolysis and activating antitumor immunity. Previously, we developed a novel OAd fully composed of human adenovirus serotype 35 (OAd35). OAd35 efficiently killed a variety of human tumor cells; however, OAd35-mediated activation of antitumor immunity remains to be evaluated. In this study, we examined whether OAd35-induced activation of immune cells contributes to the antitumor effects of OAd35.

**Methods:** Tumor infiltration and activation of immune cells following intratumoral administration of OAd35 in tumor-bearing immune-competent and nude mice were analyzed. The involvement of natural killer (NK) cells in the tumor growth-suppression effects of OAd35 was evaluated in NK cell-depleted mice. The key signals for the OAd35-mediated tumor infiltration of NK cells were examined in interferon (IFN) alpha and beta receptor subunit 1 (IFNAR1) knockout and toll-like receptor 9 (TLR9) knockout mice.

**Results:** OAd35 efficiently induced tumor infiltration of activated NK cells. NK cell depletion apparently hindered the OAd35-mediated tumor growth suppression. In IFNAR1 knockout mice, OAd35-induced tumor infiltration of activated NK cells was significantly attenuated. OAd35 did not induce tumor infiltration of NK cells in TLR9 knockout mice, although OAd35 significantly activated NK cells and showed tumor growth suppression in TLR9 knockout mice.

**Conclusions:** OAd35 significantly promoted activation and tumor infiltration of NK cells, leading to OAd35-mediated efficient tumor growth suppression. The type-I IFN signal was crucial for the OAd35-mediated tumor infiltration and activation of NK cells. The TLR9 signal was highly related to tumor infiltration of NK cells, but not NK cell activation and antitumor effects of OAd35. These findings suggest that OAd35 becomes a promising cancer immunotherapy agent *via* its enhancement of the antitumor activities of NK cells.

## BACKGROUND

Oncolytic viruses (OVs) are natural or genetically modified viruses that specifically infect and kill tumor cells without harming normal cells. Clinical trials using OVs have been conducted worldwide and have shown favorable results.^1,2^ The attractive feature of OVs is their ability to convert immunosuppressive “cold” tumors into immunologically active “hot” tumors by activating antitumor immunity. In addition to the replication-dependent direct oncolysis, OVs directly activate innate immunity in immune cells, including dendritic cells and macrophages, following phagocytosis of OVs in immune cells. Moreover, OV-mediated tumor cell lysis results in the release of not only tumor-associated antigens but also pathogen-associated molecular patterns (PAMPs), such as virus genome and damage-associated molecular patterns (DAMPs), including high mobility group box 1 (HMGB1)^3^ and heat shock proteins (HSPs),^4^ from tumor cells. The OV-induced immune responses described above lead to the activation of antitumor immunity.^5–7^ In pre-clinical and clinical trials, combination therapy of OVs and immunotherapies, such as immune checkpoint inhibitors, exhibited superior therapeutic efficacies in various types of tumors.^8,9^

Among the various types of OVs, the oncolytic adenovirus (Ad) (OAd) is the type most widely used in clinical trials of OVs.^10,11^ Tumor-selective replication of OAd is often mediated by tumor-specific expression of the Ad E1 gene, which encodes proteins essential for Ad self-replication, using a tumor cell-specific promoter.^12^ Although more than 100 types of human Ad serotypes have been identified,^13^ almost all OAds are based on human Ad serotype 5 (Ad5) (OAd5). OAd5 has various advantages as an OV; however, two major concerns of OAd5—namely, the high seroprevalence in adults^14,15^ and the low expression of the primary infection receptor, coxsackievirus-adenovirus receptor (CAR), on the malignant tumor cells,^16,17^—may limit the therapeutic efficacy of OAd5. In order to overcome these concerns regarding OAd5, we previously developed a novel OAd fully composed of human Ad serotype 35 (Ad35) (OAd35).^18^ We chose Ad35 for the development of a novel OAd because while only 20% or fewer adults have anti-Ad35 antibodies^19,20^ and the Ad35 infection receptor, human CD46, is highly expressed on a variety of human tumor cells.^21,22^ OAd35 has been shown to evade pre-existing anti-Ad5 neutralizing antibodies and to kill both CAR-positive and -negative tumor cells.^18^ Although OAd35 is a promising alternative OAd, whether OAd35 activates antitumor immunity remains to be determined. Previously, we and other groups reported that a replication-incompetent Ad35 vector more efficiently activated innate immunity in immune cells, including mouse bone marrow-derived dendritic cells, than an Ad5 vector.^23–26^ These findings led us to hypothesize that OAd35 efficiently activated innate immunity following administration, leading to efficient induction of antitumor immunity and antitumor effects.

In this study, we examined tumor infiltration and activation of immune cells following intratumoral administration of OAd35. Our results showed that OAd35 induced tumor infiltration and activation of natural killer (NK) cells more efficiently than OAd5. In addition, OAd35 mediated efficient growth suppression of subcutaneous tumors in a NK cell-dependent manner in both immune-competent and -incompetent (BALB/c nude) mice. Type-I interferon (IFN) was shown to be highly involved in both the OAd35-mediated tumor infiltration and activation of NK cells. These data indicated that NK cells play a crucial role in the antitumor effects of OAd35. OAd35 is a novel NK cell-stimulating OV and is highly promising as a cancer immunotherapy agent.

## METHODS

### Cells and viruses

H1299 (a human non-small cell lung carcinoma cell line) and B16 (a mouse skin melanoma cell line) cells were cultured in RPMI1640 (Sigma-Aldrich., St. Louis, MO) supplemented with 10% fetal bovine serum (FBS), 100 μg/ml streptomycin, and 100 U/ml penicillin in a 37 °C, 5% CO_2_ and 95% humidified air environment. Normal human peripheral blood mononuclear cells (PBMCs) (FUJIFILM Wako Pure Chemical, Osaka, Japan) were cultured in RPMI1640 (Sigma-Aldrich) supplemented with 10% FBS, 2 mM L-glutamine, 100 μg/ml streptomycin, and 100 U/ml penicillin in a 37 °C, 5% CO_2_ and 95% humidified air environment. OAd35 and OAd5 were previously produced.^18^ A replication-incompetent Ad35 vector expressing green fluorescent protein (GFP) (Ad35-GFP) was previously prepared using pAdMS18CA-GFP.^27^

### Mice

C57BL/6J mice and BALB/c nude mice (5-6 weeks, female) were purchased from Nippon SLC (Hamamatsu, Japan). IFN alpha and beta receptor subunit 1 (IFNAR1) knockout mice (5-6 weeks, C57BL/6 background) were previously obtained.^28^ Toll-like receptor 9 (TLR9) knockout mice (5-6 weeks, C57BL/6 background) were purchased from Oriental Yeast Co. (Tokyo, Japan).

### *In vitro* tumor cell lysis activities of OAds

B16 cells were seeded on a 96-well plate at 8.0×10^3^ cells/well. On the following day, cells were infected with OAd35 at 500 and 1000 virus particle (VP)/cell. Cell viabilities were measured by a WST-8 assay using a Cell Counting Kit-8 solution (Dojindo Laboratories, Kumamoto, Japan) on the indicated day points.

### *In vivo* antitumor effects of OAd35

H1299 cells (3×10^6^ cells/mouse) mixed with matrigel (Corning, Corning, NY) and B16 cells (2×10^5^ cells/mouse) were subcutaneously injected into the right flank of 5-6-week-old mice. When the tumors grew to approximately 5 to 6 mm in diameter, mice were randomly assigned into groups. OAd35 was intratumorally administrated to subcutaneous tumors at a dose of 2.0×10^9^ VP/mouse on the indicated day points. The tumor volumes were measured every three days. The following formula calculated tumor volume: tumor volume (mm^3^) = *a* x *b*^2^ x 3.14 x 6^−1^, where *a* is the longest dimension, and *b* is the shortest. For depletion of NK cells, normal rabbit serum (FUJIFILM Wako Pure Chemical) and anti-asialo GM1 antibody (FUJIFILM Wako Pure Chemical) were intravenously administered every four days from day 0. For CD8^+^ T cell depletion, 250 μg of InVivoMAb rat IgG2b isotype control antibody (clone: LTF-2) (BioXCell, Lebanon, NH) and InVivoMAb anti-mouse CD8α antibody (clone: 2.43) (BioXCell) were intraperitoneally administered every four days from day 0. For evaluation of the preventive effects of OAd35 on lung metastasis, OAd35 was intratumorally administrated to subcutaneous B16 tumors on days 0 and 3. Then, B16 cells (5×10^5^ cells per mouse) were intravenously injected into subcutaneous B16 tumor-bearing mice *via* the tail vein on day 4. Following 14 days after intravenous injection of B16 cells, mice were sacrificed, and the lung was collected. The metastatic tumor colonies on the lung were counted.

### Analysis of tumor infiltration and activation of immune cells following OAd administration

When the tumor volumes reached approximately 200 mm^3^, OAds were intratumorally administered to tumor-bearing mice at a dose of 2.0×10^9^ VP/mouse on day 0 and day 3. The blood cells in the tumor were analyzed on day 4 as follows; briefly, the tumors were surgically excised and incubated with type I collagenase (Worthington Biochemical, Lakewood, NJ). Cell numbers were counted using Cellometer Auto T4 (Nexcelom Bioscience LLC, Lawrence, MA). On day 4, cells were incubated with 7-AAD viability staining solution (Thermo Fisher Scientific, Waltham, MA), fluorescein isothiocyanate (FITC)-labeled anti-mouse CD3ε antibody (BioLegend, San Diego, CA), Pacific Blue™-labeled anti-mouse CD45 antibody (BioLegend), phycoerythrin (PE)-labeled anti-mouse CD49b (DX5) antibody (BioLegend), and allophycocyanin (APC)/cyanine7-labeled anti-mouse CD69 antibody (BioLegend) after treatment with a mouse FcR blocking reagent (Miltenyi Biotec, Bergisch Gladbach, Germany). Flow cytometric analysis was performed using a MACSQuant Analyzer (Miltenyi Biotec). The data was analyzed using FlowJo™ Software (BD Life Sciences, Ashland, OR). The gating strategy is shown in supplementary figure 1.

### Gene expression analysis in the tumors following OAd administration

OAds were intratumorally administered to tumor-bearing mice, as described above. The total RNA in the tumor was recovered using ISOGEN (Nippon Gene, Tokyo, Japan) according to the manufacturer’s instructions on day 4. cDNA was synthesized from recovered RNA by a Superscript VILO cDNA synthesis kit (Thermo Fisher Scientific). Real-time RT-PCR analysis was performed using a StepOnePlus System (Thermo Fisher Scientific) and THUNDERBIRD Next SYBR qPCR Mix reagents (TOYOBO, Osaka, Japan). The gene expression levels were normalized by the mRNA levels of a housekeeping gene, mouse glyceraldehyde-3-phosphate dehydrogenase (GAPDH). The sequences of the primers were described in supplementary table 1.

### OAd-mediated activation of human NK cells

Normal human PBMCs were seeded on a Nunclon Sphera-Treated 24-well plate (Thermo Fisher Scientific) at 1.0×10^6^ cells/well. PBMCs were infected with OAds at 100 VP/cell. After a 24-h infection, PBMCs were labeled with PE-labeled anti-human CD56 (NCAM) antibody (BioLegend), FITC-labeled anti-human CD3 antibody (BioLegend), APC/cyanine7-labeled anti-human CD69 antibody (BioLegend), and 7-AAD viability staining solution (Thermo Fisher Scientific), after treatment with a human FcR blocking reagent (Miltenyi Biotec). Flow cytometric analysis was performed as described above. The gating strategy is shown in supplementary figure 2.

### OAd genome copy numbers in the tumors

OAd35 was intratumorally administered as described above. Tumors were surgically excised at 21 days after the first administration of the anti-GM1 antibody. Total DNA in the tumors was isolated using DNAzol (Molecular Research Center, Cincinnati, OH). OAd35 genome copy numbers were quantitated by real-time PCR analysis as previously described.^18^

### Statistical analyses

Two-tailed unpaired Student’s *t*-test, Welch’s *t*-test, one-way analysis of variance (ANOVA) followed by Bonferroni’s multiple comparisons test, and two-way ANOVA with Bonferroni’s multiple comparisons test were performed using GraphPad Prism (GraphPad Software, San Diego, CA) and R (Version 4. 2. 2). Data are presented as means ± standard deviation (SD) or standard error (SE).

## Results

### Immune cell activation in the tumors following OAd35 administration

In order to examine whether OAd35 induces tumor infiltration and activation of immune cells in mice, OAd35 was intratumorally administered to B16 tumor-bearing immune-competent mice. OAd35 significantly attenuated the B16 tumor growth following intratumoral administration (figure 1A), although OAd35 did not significantly mediate *in vitro* oncolysis in B16 cells (figure 1B). Since it is well known that human Ads do not infect rodent cells,^29,30^ the *in vivo* B16 tumor growth inhibition by OAd35 did not depend on the virus infection, including virus replication, of tumor cells. These data suggested that OAd35 mediated B16 tumor growth suppression *via* immunological responses.

**Figure 1.**
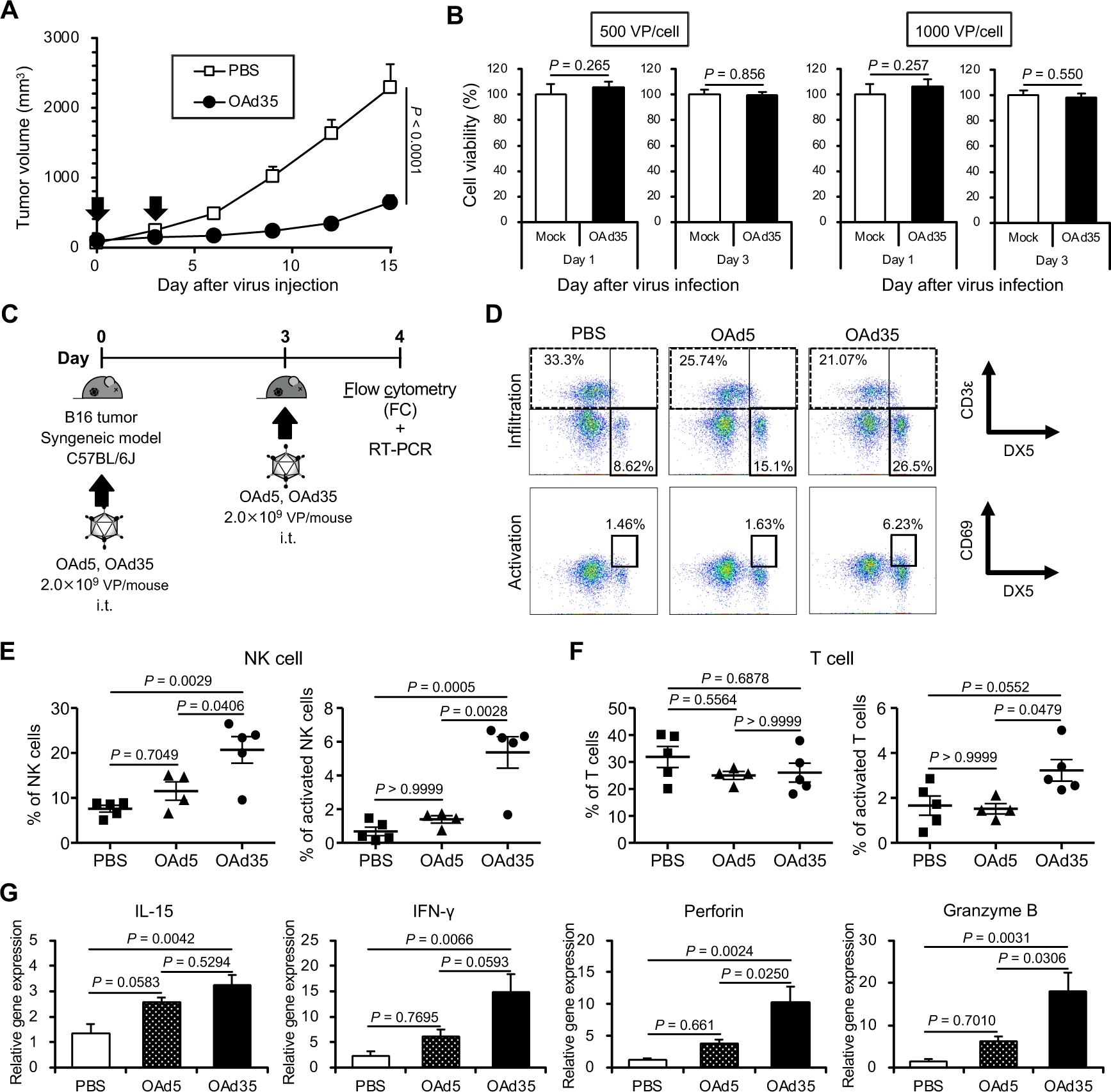
OAd35-mediated immune cell activation in C57BL/6J mice. **A)** B16 tumor growth following OAd35 administration. OAd35 was intratumorally administered at a dose of 2.0×10^9^ VP/mouse on the days indicated by arrows. Tumor volumes were expressed as the mean tumor volumes ± SE (n=8) and analyzed by two-way ANOVA with Bonferroni’s multiple comparisons test. **B)** Cell viability was determined by WST-8 assay on days 1 and 3. The viabilities of the mock-infected group were normalized to 100%. These data are expressed as the means ± SD (n=4) and analyzed by two-tailed unpaired Student’s *t*-test. **C)** Experimental scheme for the analysis of tumor infiltration of immune cells following intratumoral injection of OAds in C57BL/6J mice bearing B16 tumors. **D)** Representative dot plots of tumor-infiltrated NK cells (CD3ε^−^/DX5^+^ cells) and T cells (CD3ε^+^ cells) (left panel) and CD69^+^/CD3ε^−^/DX5^+^ cells (right panel) following OAds administration. **E)** The percentages of tumor-infiltrated NK cells and activated NK cells and **F)** tumor-infiltrated T cells and activated T cells on day 4. The data were expressed as the means ± SE (PBS, OAd35: n=5; OAd5: n=4). Data represent two independent experiments. **G)** The mRNA levels of IL-15, IFN-γ, perforin, and granzyme B in the tumors following OAds administration. The gene expression levels were normalized by GAPDH. These data were expressed as the means ± SE (n=5). One-way ANOVA with Bonferroni’s multiple comparisons post hoc test was performed.

Next, we examined the tumor infiltration and activation of immune cells following intratumoral administration of OAd35 (figure 1C). OAd5 was also intratumorally injected as a control. The percentages of CD3ε^−^/DX5^+^ cells (total NK cells) and CD69^+^/DX5^+^ cells (activated NK cells) were significantly increased by intratumoral injection of OAd35 on day 4 (figure 1D, E). While CD69 expression on CD3ε^+^ T cells was significantly higher following administration of OAd35 than following administration of OAd5, the percentages of CD3ε^+^ T cells in the tumors were not increased on day 4. OAd5 less efficiently induced activation and tumor infiltration of NK cells in B16 tumor-bearing mice than OAd35 on day 4. In addition, approximately 4-fold higher levels of CD69^+^ human NK cells in human PBMCs were found following incubation of human PBMCs with OAd35, compared with incubation with OAd5 (supplementary figure 3). The mRNA expressions of IFN-γ, perforin, and granzyme B, which were highly expressed in activated NK cells,^31^ and interleukin (IL)-15 in the tumors were significantly increased by intratumoral administration of OAd35 (figure 1F). These results indicated that OAd35 enhanced the tumor infiltration of activated NK cells and significantly inhibited tumor growth in a virus infection-independent manner.

### Attenuation of OAd35-mediated tumor growth suppression in NK cell-depleted mice

In order to examine the involvement of NK cells in OAd35-mediated tumor growth suppression in immune-competent mice, anti-GM1 antibodies were intravenously administered before intratumoral administration of OAd35 to remove NK cells from the mice (figure 2A). In PBS-injected groups, no apparent differences in B16 tumor growth were observed between the naïve serum-pretreated and anti-GM1 antibody-pretreated groups (figure 2B). OAd35 significantly suppressed the B16 tumor growth of naïve serum-pretreated mice; on the other hand, when NK cells were depleted by anti-GM1 antibodies, the OAd35-mediated suppression of B16 tumor growth disappeared.

**Figure 2.**
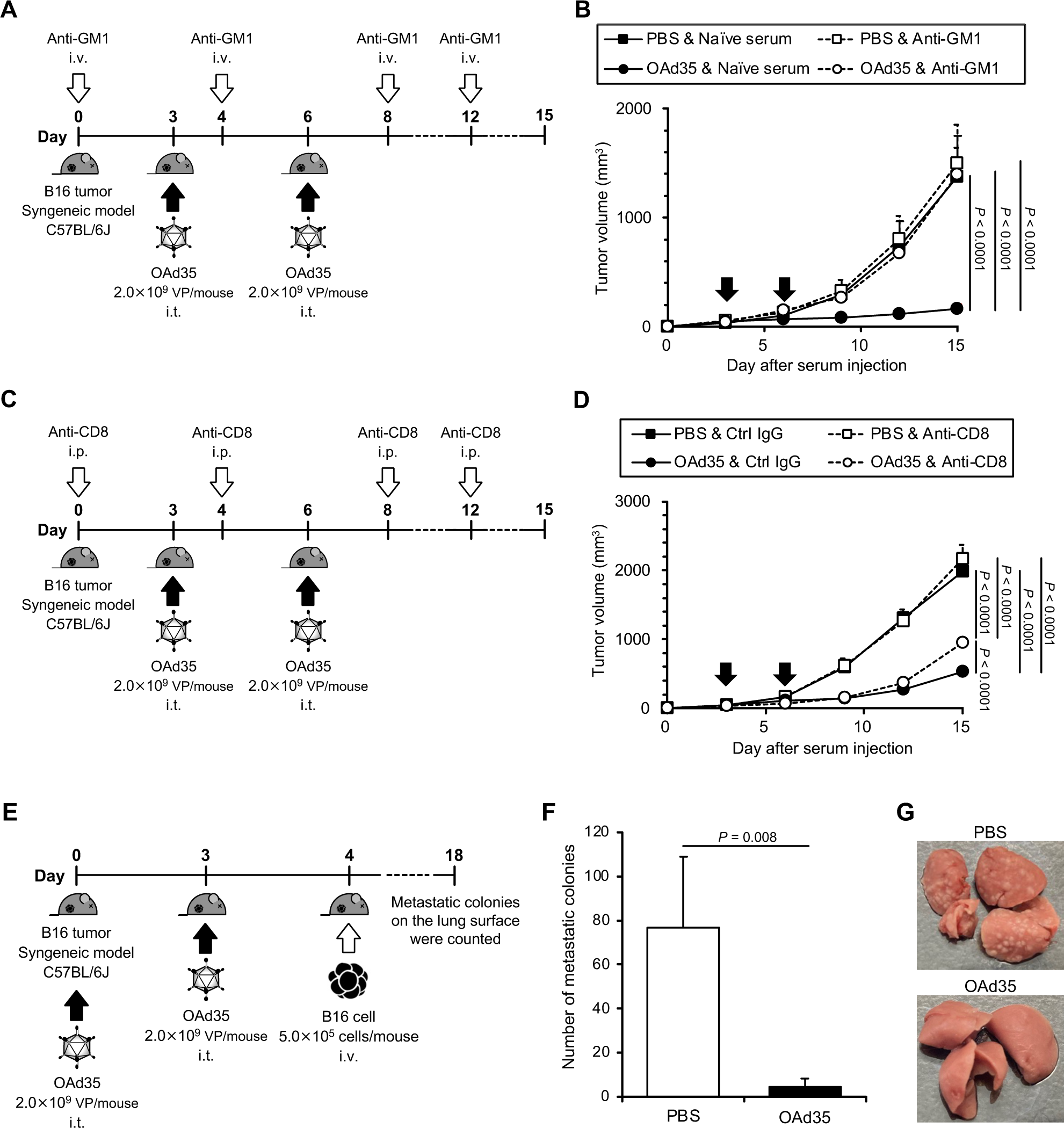
The tumor growth suppression levels of OAd35 in NK cell-depleted C57BL/6J mice. **A)** Experimental strategy for analysis of the antitumor effects of OAd35 in NK cell-depleted C57BL/6J mice bearing B16 tumor. **B)** B16 tumor growth following OAd35 administration in naïve serum-pretreated and anti-GM1 antibody-pretreated mice. The arrows indicate the days of OAd35 injection. The tumor volumes were expressed as the means ± SE (PBS: n=6; OAd35: n=8). Two-way ANOVA followed by Bonferroni’s multiple comparisons test was performed. **C)** Experimental scheme for analysis of the involvement of CD8^+^ T cells on the tumor growth suppression effects of OAd35. **D)** B16 tumor growth following OAd35 administration in control IgG-preinjected and anti-CD8α antibody-preinjected mice. The black arrows indicate the day points of OAd35 injection. The tumor volumes were expressed as the means ± SE (PBS: n=6; OAd35: n=8). Two-way ANOVA followed by Bonferroni’s multiple comparisons test was performed. **E)** Experimental design to analyze the suppressive effects of intratumorally administered OAd35 on B16 lung metastasis. **F)** The numbers of B16 metastatic colonies on the lung surface following OAd35 administration with or without pretreatment with anti-GM1 antibody. These data were expressed as the means ± SE (PBS: n=6-8; OAd35: n=6). One-way ANOVA with Bonferroni’s multiple comparisons post hoc test was performed. **G)** Representative images of the lungs from the PBS and OAd35 administration groups.

To evaluate the involvement of CD8^+^ T cells in the antitumor effects of OAd35, CD8^+^ T cells in B16 tumor-bearing mice were depleted by intraperitoneal injection of anti-CD8α antibody, followed by intratumoral administration of OAd35 (figure 2C). Comparable volumes of B16 tumors were found following OAd35 administration in the mice receiving anti-CD8α antibody and control IgG from day 3 to day 12 (figure 2D). CD8^+^ T cell depletion resulted in slight but statistically significant inhibition of the tumor growth suppression on day 15 following OAd35 administration. These results indicated that CD8^+^ T cells slightly but significantly contributed to OAd35-mediated tumor growth suppression in B16 tumors, especially at late time points.

Next, in order to evaluate the preventive effects of OAd35 on lung metastasis, B16 cells were intravenously injected one day after intratumoral administration of OAd35 (figure 2E). NK cells were highly involved in the inhibition of lung metastasis of B16 cells.^32,33^ The number of metastatic colonies in the lung was significantly decreased in OAd35-treated mice (figure 2F, G). These results indicated that NK cells played a key role in the antitumor effects of OAd35 in B16 tumor-bearing mice.

### OAd35-induced tumor infiltration of NK cells in human tumor-bearing nude mice

In order to examine whether OAd35 mediates tumor growth suppression *via* NK cell activation even in nude mice, OAd35 was intratumorally administrated to H1299 tumor-bearing nude mice (figure 3A). BALB/c nude mice possess functional NK cells at levels comparable to or higher than those in wild-type BALB/c mice.^34,35^ Intratumoral administration of OAd35 significantly increased the percentages of CD3ε^−^/DX5^+^ cells (total NK cells) and CD69^+^/DX5^+^ cells (activated NK cells) in H1299 tumors (figure 3B, C). The OAd35-mediated activation and tumor infiltration levels of mouse NK cells were significantly higher than those by conventional OAd5. OAd35 induced a significant and efficient increase in the mRNA levels of IL-15, IFN-γ, perforin, and granzyme B compared to those in the other groups (figure 3D). These results indicated that OAd35 more efficiently induced activation and tumor infiltration of NK cells in nude mice than OAd5.

**Figure 3.**
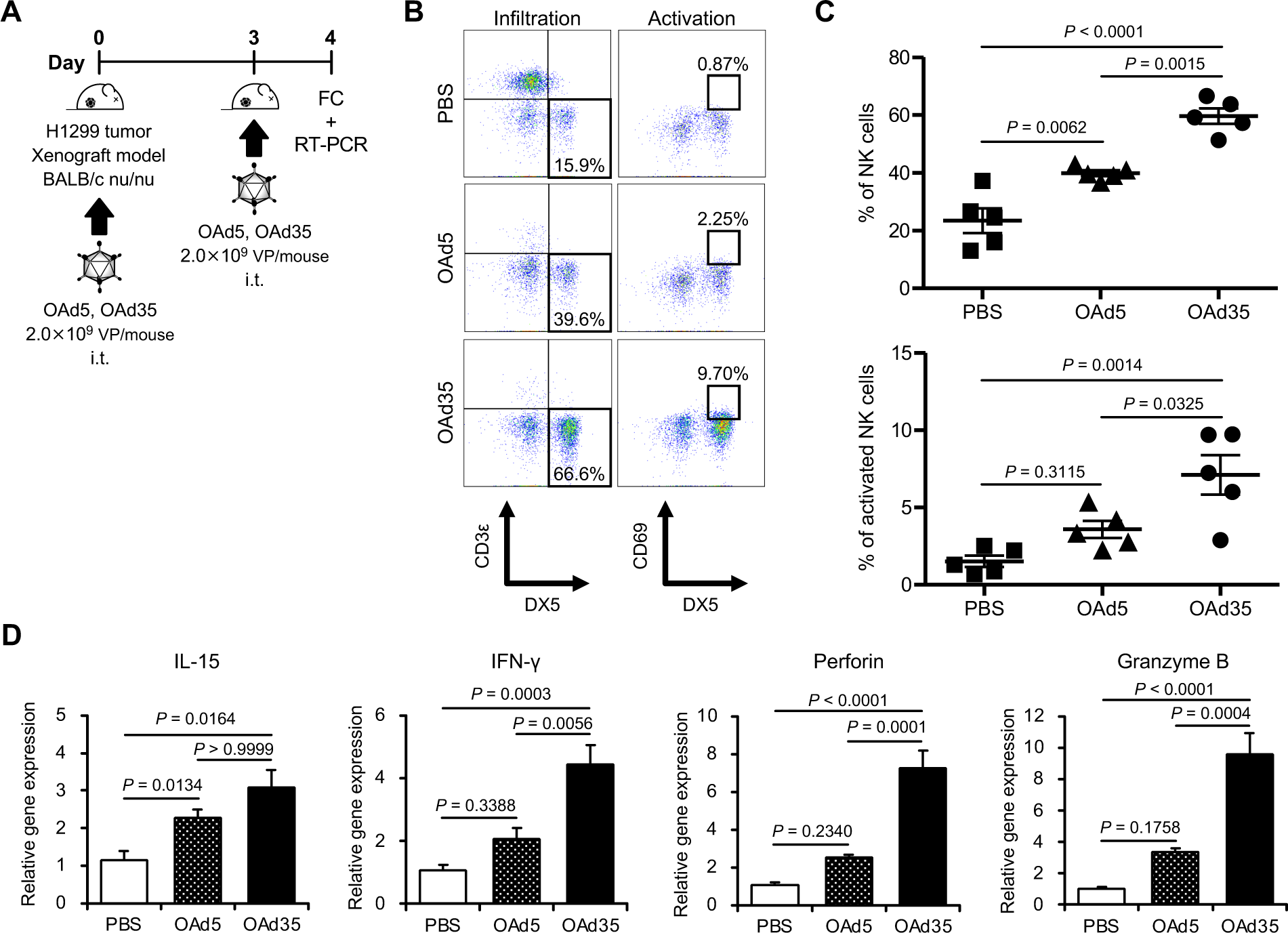
Tumor infiltration of activated NK cells after OAd35 administration in BALB/c nude mice. **A)** Experimental design for analysis of the tumor infiltration of activated NK cells following intratumoral injection of OAds in BALB/c nude mice bearing H1299 tumors. **B)** Representative dot plots of CD3ε^−^/DX5^+^ cells in CD45^+^ cells (left panel) and CD69^+^/DX5^+^ cells in CD3ε^−^/CD45^+^ cells (right panel). **C)** The percentages of CD3ε^−^/DX5^+^ cells in CD45^+^ cells (upper panel) and of CD69^+^/DX5^+^ cells in CD3ε^−^/CD45^+^ cells (lower panel) on day 4. Data represent two independent experiments. **D)** The mRNA expression levels in the tumors following OAds administration. The gene expression levels were normalized by GAPDH. These data were expressed as the means ± SE (n=5). One-way ANOVA with Bonferroni’s multiple comparisons post hoc test was performed.

### Impact of NK cells on the antitumor effects and replication of OAd35 in nude mice

In order to examine the impact of NK cells on the antitumor effects and replication of OAd35, NK cells in nude mice were depleted by intravenous injection of anti-GM1 antibodies, followed by intratumoral administration of OAd35 (figure 4A). In the presence of NK cells, intratumoral administration of OAd35 resulted in significant suppression of the subcutaneous H1299 tumor growth (figure 4B). Conversely, NK cell depletion clearly canceled the tumor growth inhibition of OAd35. These results indicated that NK cells filled an essential role in the tumor growth suppression effects of OAd35 in immune-incompetent mice.

**Figure 4.**
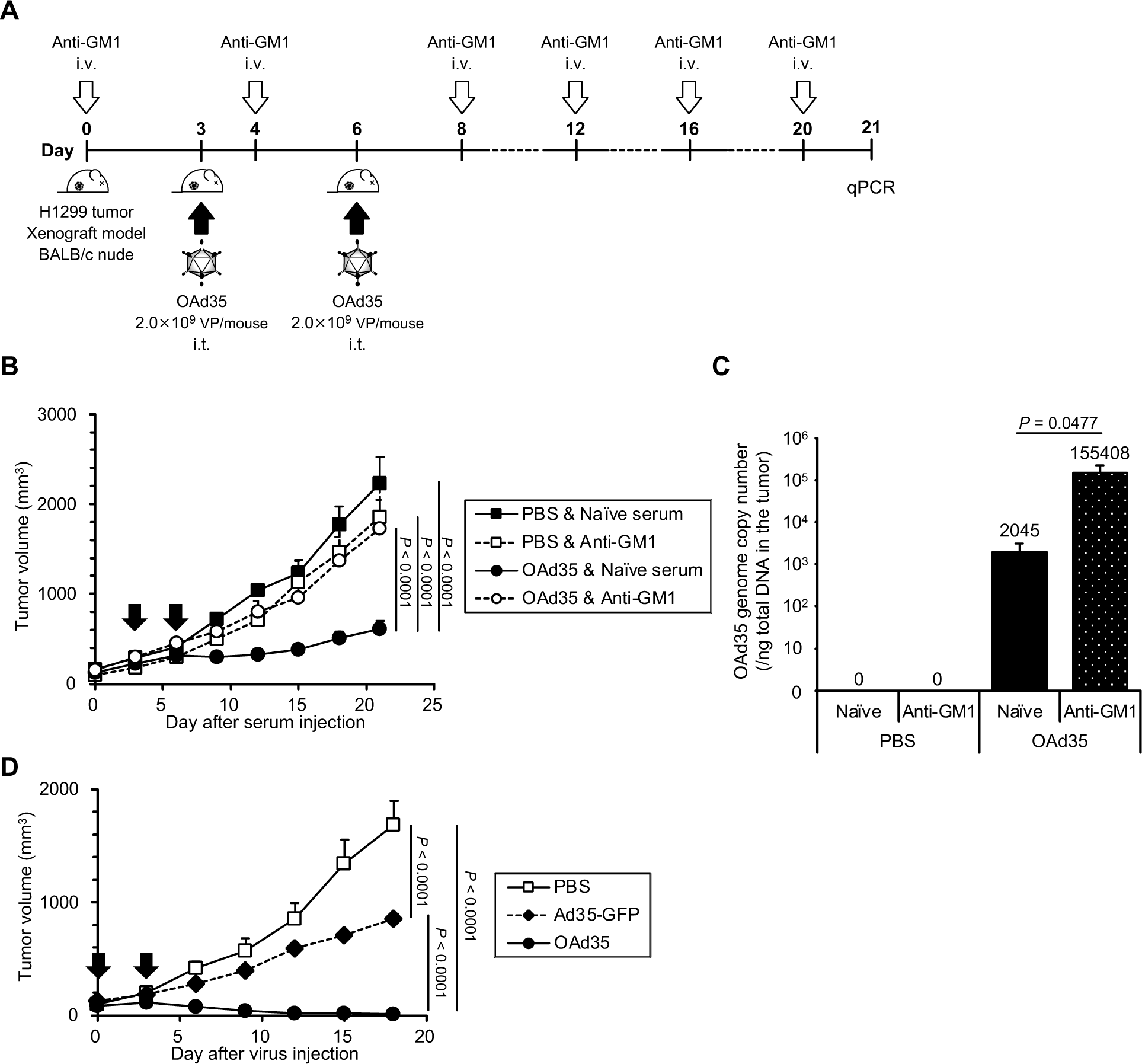
Impact of NK cells on the antitumor effects of OAd35 in BALB/c nude mice. **A)** Experimental schedule for analysis of the involvement of NK cells in the H1299 tumor growth suppression levels by intratumorally administered OAd35 in BALB/c nude mice. **B)** H1299 tumor growth following OAd35 administration. The arrows indicate the days of the OAd35 injection. These data were expressed as the means ± SE (PBS: n=5; OAd35: n=7). **C)** The OAd35 genome copy numbers in the tumors following OAd35 administration. The OAd35 genome copy numbers in the tumor were determined by quantitative PCR analysis at day 21. These data were expressed as the means ± SE (PBS: n=5; OAd35: n=7). One-way ANOVA followed by Bonferroni’s multiple comparisons test was performed. **D)** H1299 tumor growth following intratumoral administration of Ad35-GFP and OAd35. Replication-incompetent Ad35-GFP and OAd35 were intratumorally administered to BALB/c nude mice bearing H1299 tumors at a dose of 2.0×10^9^ VP/mouse on the days indicated by arrows. Tumor volumes were expressed as the mean tumor volumes ± SE (n=7) and analyzed by two-way ANOVA with Bonferroni’s multiple comparisons post hoc test.

Next, in order to examine the virus replication levels in the tumors, the OAd35 genome copy numbers in the tumor were measured by quantitative PCR analysis at 21 days after the first administration of anti-GM1 antibodies (figure 4C). The results showed that NK cell depletion significantly increased the OAd35 genome copy numbers in H1299 tumors by approximately 80-fold compared to those in the tumors of naïve serum-treated mice.

In order to further examine whether OAd35 replication in the tumors is crucial for OAd35-mediated tumor growth suppression, a replication-incompetent Ad35-GFP was intratumorally administered. OAd35 more efficiently suppressed the tumor growth than Ad35-GFP (figure 4D), indicating that OAd35 infection in H1299 tumors was important for the OAd35-mediated growth suppression of subcutaneous H1299 tumors following intratumoral administration.

### Contribution of the type-I IFN signal to OAd35-mediated NK cell activation and tumor growth inhibition

To examine the contribution of the type-I IFN signal to OAd35-mediated tumor infiltration of activated NK cells and tumor growth suppression, OAd35 was intratumorally administered to IFNAR1 knockout mice bearing B16 tumors (figure 5A). In IFNAR1 knockout mice, there was no apparent tumor infiltration of NK cells following intratumoral administration of OAd35 (figure 5B, C). The CD69 expression levels on NK cells in the B16 tumors were comparable in PBS- and OAd35-administered IFNAR1 knockout mice. The mRNA expression levels of IL-15, IFN-γ, perforin, and granzyme B in the tumors were not significantly increased following OAd35 administration in IFNAR1 knockout mice (figure 5D). No growth suppression of B16 tumors was apparent following OAd35 administration in IFNAR1 knockout mice (figure 5E). These results indicated that OAd35 recruited activated NK cells into the tumors and mediated tumor growth suppression in a type-I IFN-dependent manner.

**Figure 5.**
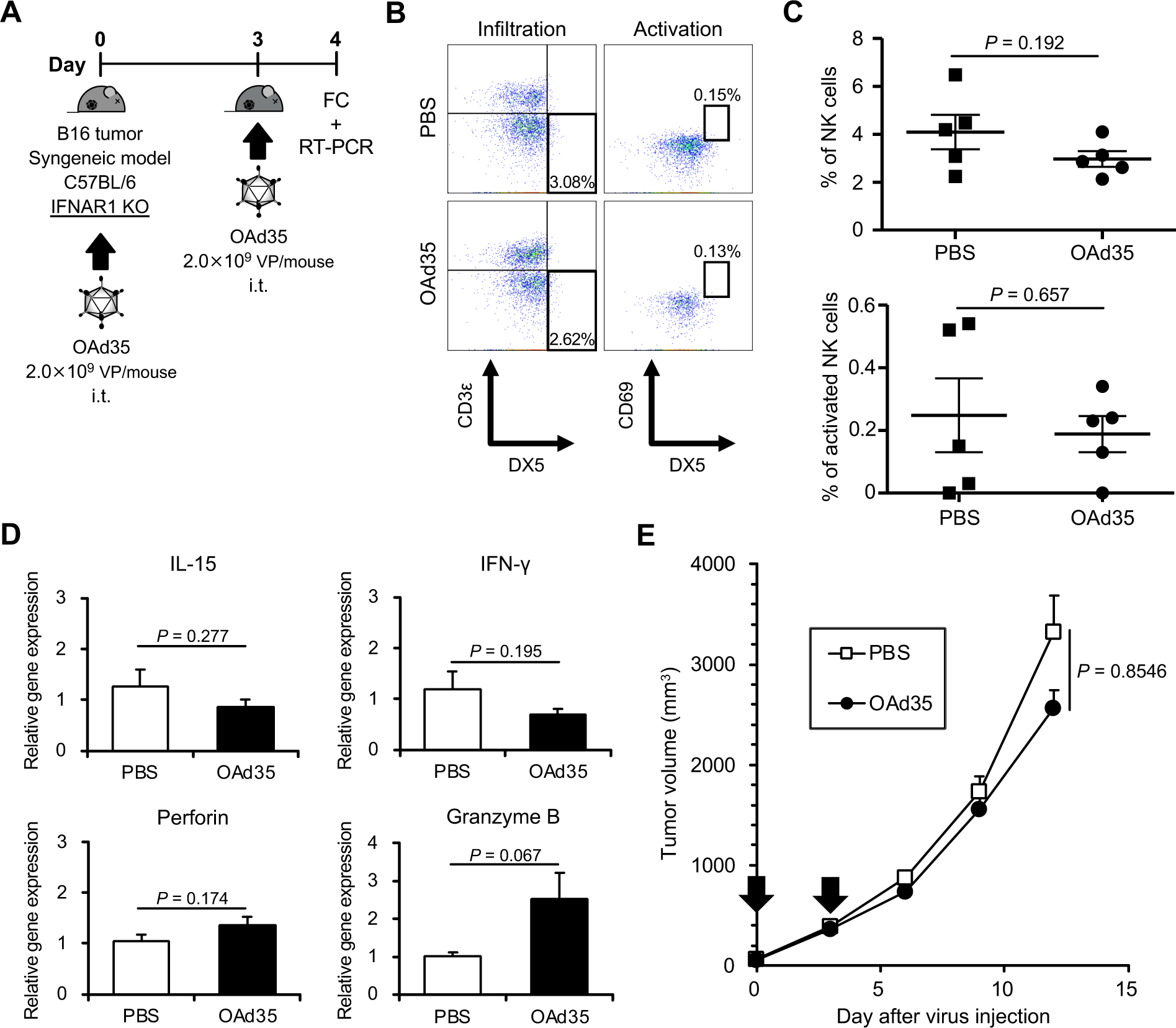
The role of the type-I IFN signal on OAd35-mediated tumor infiltration of NK cells and antitumor effects. **A)** Experimental strategy for analysis of the involvement of the type-I IFN signal in NK cell infiltration following intratumoral injection of OAd35 in IFNAR1 knockout mice. **B)** Representative dot plots of CD3ε^−^/DX5^+^ cells in CD45^+^ cells (left panel) and CD69^+^/DX5^+^ cells in CD3ε^−^/CD45^+^ cells (right panel) in the tumors. **C)** The percentages of CD3ε^−^/DX5^+^ cells in CD45^+^ cells (upper panel) and CD69^+^/DX5^+^ cells in CD3ε^−^/CD45^+^ cells (lower panel) in the tumors on day 4. Data represent two independent experiments. **D)** The mRNA expression levels in the tumors following OAd35 administration. The gene expression levels were normalized by GAPDH. These data were expressed as the means ± SE (n=5). Data were analyzed by two-tailed unpaired Student’s *t*-test. **E)** OAd35 was intratumorally administered at a dose of 2.0×10^9^ VP/mouse on the days indicated by arrows. Tumor volumes were expressed as the mean tumor volumes ± SE (n=8) and analyzed by two-way ANOVA followed by Bonferroni’s multiple comparisons test.

### Involvement of the TLR9 signal in OAd35-mediated tumor infiltration of NK cells

To examine the involvement of the TLR9 signal in OAd35-mediated activation and tumor infiltration of NK cells, OAd35 was intratumorally administered to TLR9 knockout mice bearing B16 tumors (figure 6A). Previous studies demonstrated that TLR9 was involved in Ad35-induced innate immune responses.^24^ The percentages of CD3ε^−^/DX5^+^ cells (total NK cells) were not significantly increased by intratumoral injection of OAd35 in TLR9 knockout mice, although the percentages of CD69^+^/DX5^+^ cells in OAd35-treated mice were significantly higher than those in PBS-treated mice (figure 6B, C). Significant elevations in the mRNA expression levels of IL-15, IFN-γ, perforin, and granzyme B in the tumors were found following OAd35 administration in TLR9 knockout mice (figure 6D). While OAd35 did not induce NK cell infiltration, intratumoral administration of OAd35 significantly suppressed B16 tumor growth in TLR9 KO mice (figure 6E). These results indicated that the TLR9 signal was involved in the OAd35-mediated tumor infiltration of NK cells, but not in the activation and antitumor effects of OAd35.

**Figure 6.**
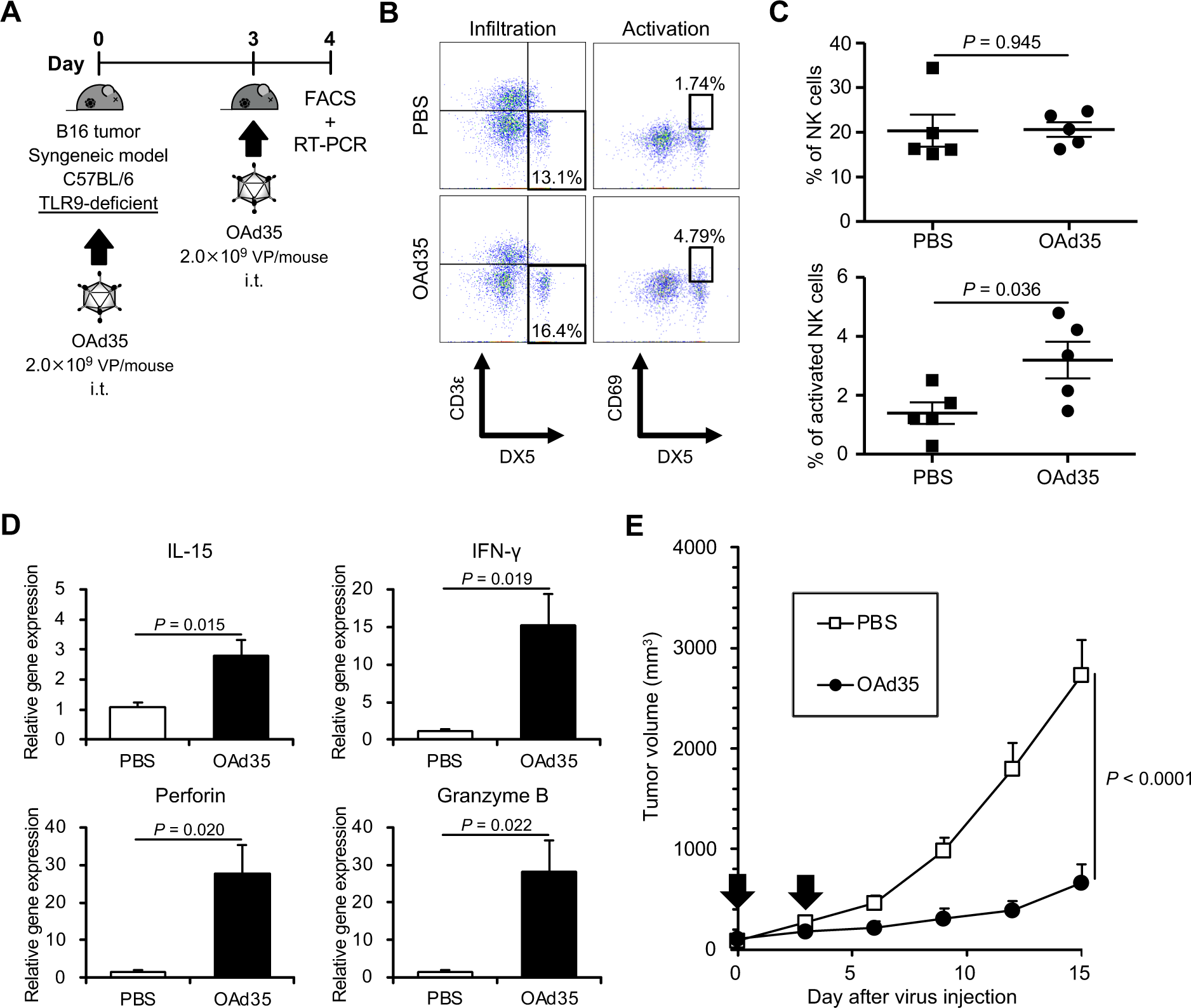
Impact of TLR9 on the tumor infiltration of activated NK cells following OAd35 administration. **A)** Experimental design for analysis of the involvement of TLR9 in OAd35-mediated tumor infiltration of NK cells using TLR9 knockout mice. **B)** Representative dot plots of CD3ε^−^/DX5^+^ cells in CD45^+^ cells (left panel) and CD69^+^/DX5^+^ cells in CD3ε^−^/CD45^+^ cells (right panel) in the tumors following OAd35 administration. **C)** The percentages of CD3ε^−^/DX5^+^ cells in CD45^+^ cells (upper panel) and CD69^+^/DX5^+^ cells in CD3ε^−^/CD45^+^ cells (lower panel) in the tumors. These data were expressed as the means ± SE (n=5) and analyzed by two-tailed unpaired Student’s t-test. Data represent two independent experiments. **D)** The mRNA expression levels in the tumors following OAd35 administration. The gene expression levels were normalized by GAPDH. These data were expressed as the means ± SE (n=6). Data were analyzed by two-tailed two-tailed Welch’s t-test. **E)** OAd35 was intratumorally administered at a dose of 2.0×10^9^ VP/mouse on the days indicated by arrows. Tumor volumes were expressed as the mean tumor volumes ± SE (PBS: n=7; OAd35: n=8) and analyzed by two-way ANOVA with Bonferroni’s multiple comparisons test.

## Discussion

The aim of this study was to reveal the mechanisms of OAd35-mediated tumor growth suppression following intratumoral administration in subcutaneous tumor-bearing mice. The results showed that NK cells were efficiently activated and recruited to tumors following OAd35 administration in both immune-competent and nude mice (figure 1C, E, 3B, C). OAd5 mediated significantly lower levels of tumor infiltration of NK cells than OAd35. Moreover, NK cells largely contributed to OAd35-mediated tumor growth suppression (figure 2B, 4B). It is well known that NK cells play a crucial role in antitumor and antivirus responses. Previous studies reported that tumor infiltration and activation levels of NK cells were highly related to cancer metastasis, malignancy, and development rates in mouse models and patient cohorts.^36–40^ In addition, tumor infiltration levels of NK cells correlated well with the survival rates of head and neck cancer patients.^41^ Hence, NK cells have been gaining much attention as a key player in cancer immunotherapy.^42,43^ Promotion of tumor infiltration and activation of NK cells are both highly important for OV-mediated cancer therapy.^44^ Other OVs, including reovirus^45^ and herpesvirus,^46^ also enhanced the tumor infiltration and activation of NK cells following administration and showed efficient antitumor effects in clinical trials.

In our present experiments, OAd35 did not appear to induce NK cell recruitment into the tumor in IFNAR1 knockout mice or TLR9 knockout mice (figure 5, 6). These data indicated that TLR9 and type-I IFN signaling were crucial for the OAd35-mediated tumor infiltration of NK cells. Based on all the above, we can now consider the following hypothesis in regard to the mechanism of OAd35-mediated NK cell recruitment. First, OAd35 induced type-I IFN production in immune cells, including dendritic cells and macrophages, in a TLR9-dependent manner following administration. Previous studies demonstrated that wild-type Ad35 and a replication-incompetent Ad35 vector more efficiently induced type-I IFN production in dendritic cells than wild-type Ad5 and a replication-incompetent Ad5 vector.^24–26^ Next, type-I IFN efficiently activated NK cells. Type-I IFN is known to be a potent activator of NK cells. The activated NK cells, in turn, efficiently killed tumor cells.

In TLR9 knockout mice, intratumoral administration of OAd35 did not induce tumor infiltration of NK cells; however, it activated NK cells and showed significant growth suppression (figure 6). There are two possibilities. First, other DNA receptors, such as cyclic GMP-AMP synthase (cGAS), may have recognized the OAd35 genome and activated NK cells in a type I IFN-dependent manner. OAds in this study modified the E1A gene to circumvent the binding of the E1A protein and stimulation of IFN genes (STING).^18^ Thus the cGAS-STING pathway or other signals might contribute to the activation of NK cells in TLR9 knockout mice. Second, NK cells directly interacted with OAd35-infected cells. It is possible that the upregulation of NK cell activation ligands or the downregulation of inhibition ligands would be induced on OAd35-infected cells. Further studies are required to reveal all contributors to the OAd35-mediated NK cell activation.

There was no apparent recruitment of the activated NK cells in B16 and H1299 tumors following OAd5 administration (figure 1E, 3C). In addition, the mRNA levels of IFN-γ, perforin and granzyme B in the tumors were not significantly elevated following OAd5 administration in tumor-bearing mice (figure 1G, 3D). These data suggested that OAd5 less efficiently activated innate immunity following administration compared to OAd35. Wild-type Ad35 and a replication-incompetent Ad35 vector induced higher amounts of type-I IFN production than wild-type Ad5 and a replication-incompetent Ad5 vector in cultured dendritic cells.^25^ We consider that the differences in the activation levels of innate immunity between Ad5 and Ad35 were partially due to the differences in intracellular trafficking. Because the amounts of Ad35 enclosed in late endosomes were higher than the corresponding amounts of Ad5,^47^ Ad35 had more chances to interact with TLR9 in the endosomes. Further investigations into the differences among these immune profiles will contribute to the future design of immune stimulating medicines.

Although NK cell depletion induced by anti-GM1 antibody in nude mice significantly increased the OAd35 genome copy numbers in the tumors, OAd35 did not suppress the H1299 tumor growth in the absence of NK cells (figure 4B, 4C). We previously demonstrated that OAd35 efficiently replicated in H1299 cells *in vitro*, leading to efficient lysis of H1299 cells following infection.^18^ Probably, OAd35-mediated activation of antitumor immunity, including activation of NK cells, made a greater contribution to the tumor growth suppression following administration than virus replication-mediated tumor cell lysis. In order to achieve efficient therapeutic effects of OAd35 in cancer patients, it is important to examine whether NK cells are efficiently activated by OAd35 in cancer patients rather than whether OAd35 efficiently replicates in the cancer cells of patients.

The contribution of CD8^+^ T cells in the OAd35-mediated tumor growth suppression was clearly weaker than that of NK cells (figure 2B, D). OAd35 genome copy numbers in the tumors were significantly restored by treatment with anti-GM1 antibody, suggesting that OAd35-mediated NK cell activation led to the clearance of OAd35-infected tumor cells (figure 4C). The significant clearance of OAd35-infected tumor cells by NK cells might relate to the lesser contribution of CD8^+^ T cells to the OAd35-mediated antitumor effects. Cytotoxic CD8^+^ T cells play a huge role in antitumor immunity, including oncolytic virus-induced antitumor immunity. In order to maximize the therapeutic efficacy of OAd35, further study is needed to understand the relationship between OAd35 and cytotoxic CD8^+^ T cells.

Antitumor therapeutic genes are often incorporated into the OV genome (so-called “arming”) to enhance the antitumor effects of OVs. Incorporation of immunostimulatory genes, such as IL-2,^48^ IL-12,^49^ IL-15,^50^ and full-length antibody,^51^ into OV genomes has been shown to significantly enhance the tumor infiltration of NK cells, leading to efficient tumor growth suppression. These genetic modifications of the OAd35 genome would enhance the antitumor effects of OAd35 by inducing further activation and tumor infiltration of NK cells.

Since human Ads cannot replicate in rodent cells, the antitumor effects of OAds are commonly assessed in human tumor-bearing immune-deficient mice, such as nude mice, severe combined immune deficiency (SCID) mice, or nonobese diabetic (NOD)-SCID mice. Although innate immunity works in these immune-deficient mice, it is often overlooked.^34,35^ Our findings indicated that NK cells in BALB/c nude mice were significantly activated *via* OAd35-induced innate immunity and contributed to the tumor growth inhibition of OAd35 (figure 4B). Nude mice possess various types of immune cells other than T cells. When the antitumor effects of OVs are assessed in nude mice, we should pay attention to the involvement of NK cells and other innate immune cells in the antitumor effects.

In conclusion, we revealed that OAd35 efficiently suppressed subcutaneous tumor growth in a virus infection-independent and NK cell-dependent manner following intratumoral administration. Type-I IFN signaling was involved in OAd35-mediated tumor infiltration and activation. These findings are essential for the clinical investigation of OAd35, and OAd35 is expected to become a promising cancer immunotherapy agent that enhances the antitumor activities of NK cells.

## Declarations

### Ethics approval

The Animal Experiment Committee of Osaka University approved the animal experiments.

### Consent for publication

All authors have read and approved the submission of the manuscript.

### Availability of data and material

All data generated or analyzed during this study are included in this article and supplemental information.

### Authors’ contributions

**Ryosuke Ono:** Conceptualization; Data curation; Formal analysis; Funding acquisition; Project administration; Investigation; Methodology; Validation; Visualization; Writing - original draft. **Fuminori Sakurai:** Funding acquisition; Project administration; Supervision; Writing - review & editing. **Ken J. Ishii:** Resources. **Hiroyuki Mizuguchi:** Supervision; Funding acquisition; Writing - review & editing.

## Acknowledgments

We thank Kosuke Takayama (Graduate School of Pharmaceutical Sciences, Osaka University, Osaka, Japan) and Kazunori Aoki (National Cancer Center Research Institute, Tokyo, Japan) for their support.

## List of Abbreviations

Ad: Adenovirus
Ad5: Adenovirus serotype 5
Ad35: Adenovirus serotype 35
APC: Allophycocyanin
CAR: Coxsackievirus-adenovirus receptor
cGAS: Cyclic GMP-AMP synthase
DAMPs: Damage-associated molecular patterns
FITC: Fluorescein isothiocyanate
HSPs: Heat shock proteins
HMGB1: High mobility group box 1
IFN: Interferon
IFNAR1: Interferon alpha and beta receptor subunit 1
IL: Interleukin
NK cell: Natural killer cell
NOD: Nonobese diabetic
Oad: Oncolytic adenovirus
OAd5: Oncolytic adenovirus serotype 5
OAd35: Oncolytic adenovirus serotype 35
Ovs: Oncolytic viruses
PAMPs: Pathogen-associated molecular patterns
PBMCs: Peripheral blood mononuclear cells
PE: Phycoerythrin
SCID: Severe combined immune deficiency
STING: Stimulation of IFN genes
TLR9: Toll-like receptor 9
VP: Virus particles

---

**Figure S1.**
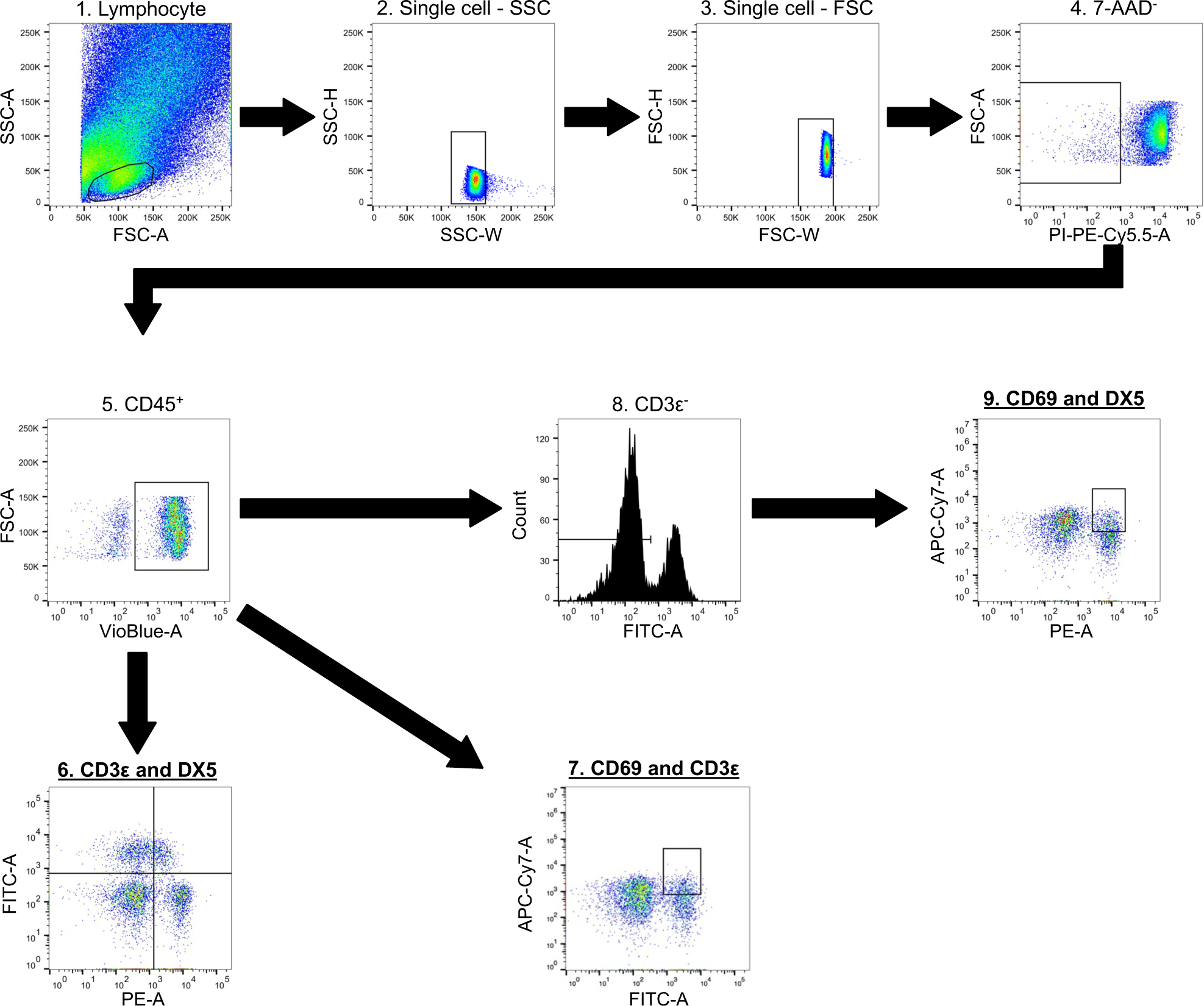
The gating strategy used in analysis of tumor-infiltrated and -activated immune cells. Cells were isolated from the B16 tumor on the C57BL/6J mice. Single lymphocytes were gated on forward scatter (FSC) and side scatter (SSC). 7AAD^−^ and CD45^+^ living lymphocytes were then further gated to determine CD3ε^−^/DX5^+^ (NK cells), CD3ε^+^/DX5^−^ (T cells), CD3ε^−^/DX5^+^/CD69^+^ (Activated NK cells), and CD3ε^+^/CD69^+^ (Activated T cells).

**Figure S2.**
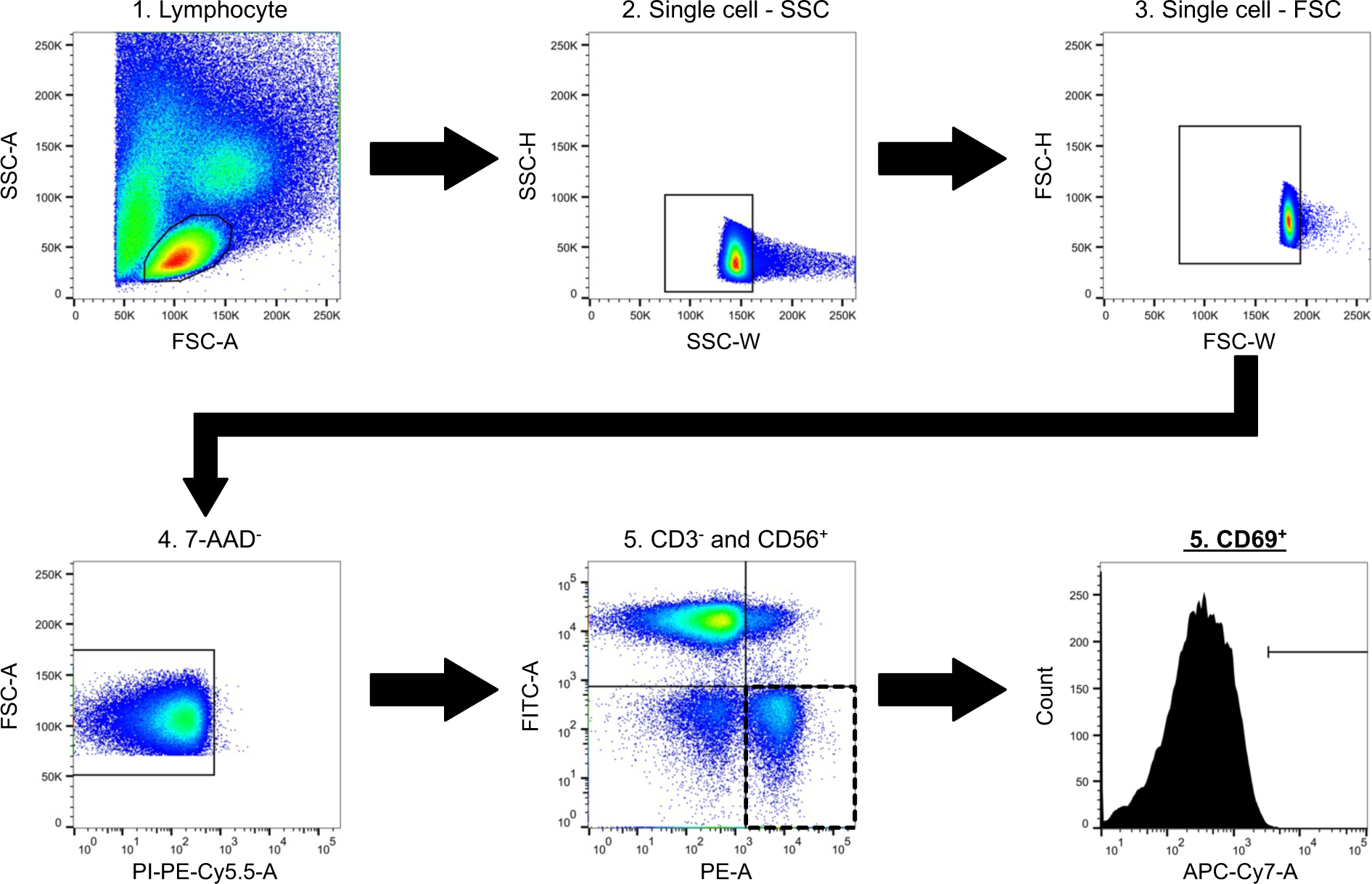
The gating strategy for analysis of human PBMCs. Single lymphocytes in PBMCs were gated on FSC and SSC. 7AAD^−^ living cells were gated by CD3^−^/CD56^+^ (NK cells) and then the expression of CD69 (Activated NK cells) was determined.

**Figure S3.**
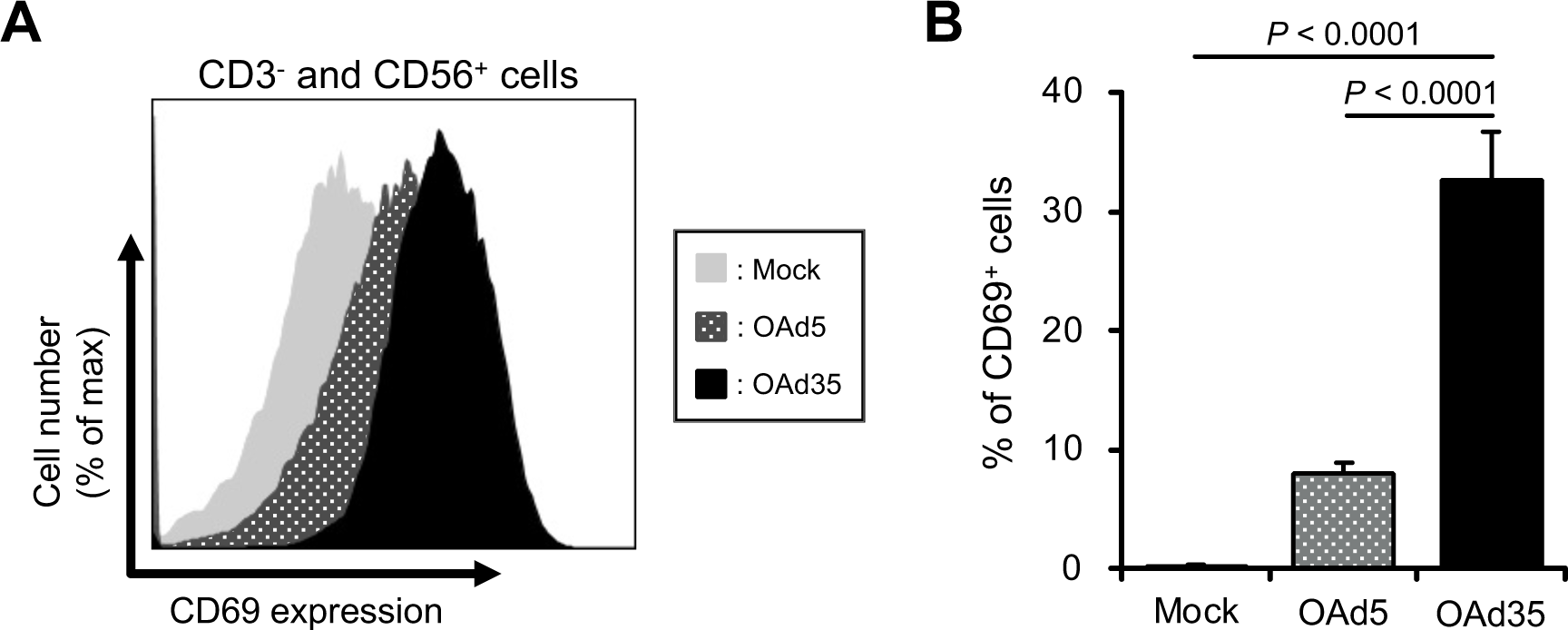
The OAd35-mediated activation of NK cells in human PBMCs. **A)** Human PBMCs were incubated with OAds at 100 VP/cell. Following 24 h of incubation with a shake, the CD69 expression in the CD3^−^ and CD56^+^ population was measured by flow cytometry. These histograms represented quadruplicate experiments. **B)** The percentages of CD69^+^ cells in the CD3^−^ and CD56^+^ cells. These were expressed as the means ± SD (n=4). One-way ANOVA followed by Dunnett’s multiple comparisons test was performed (vs. OAd35).

**Table S1.**
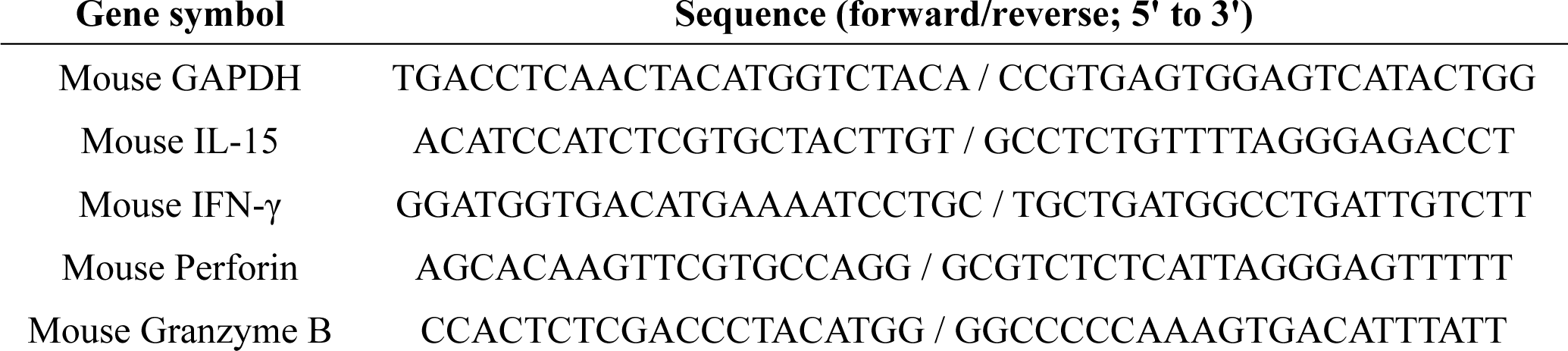
Primer sequences.

## Reference

1. Raja J, Ludwig JM, Gettinger SN, Schalper KA, Kim HS. Oncolytic virus immunotherapy: future prospects for oncology. J Immunother Cancer. 2018;6(1):140.

2. Hemminki O, Dos Santos JM, Hemminki A. Oncolytic viruses for cancer immunotherapy. J Hematol Oncol. 2020;13(1):84.

3. Huang B, Sikorski R, Kirn DH, Thorne SH. Synergistic anti-tumor effects between oncolytic vaccinia virus and paclitaxel are mediated by the IFN response and HMGB1. Gene Ther. 2011;18(2):164–172.

4. Moehler M, Zeidler M, Schede J, et al. Oncolytic parvovirus H1 induces release of heat-shock protein HSP72 in susceptible human tumor cells but may not affect primary immune cells. Cancer Gene Ther. 2003;10(6):477–480.

5. Kaufman HL, Kohlhapp FJ, Zloza A. Oncolytic viruses: a new class of immunotherapy drugs. Nat Rev Drug Discov. 2015;14(9):642–662.

6. Bommareddy PK, Shettigar M, Kaufman HL. Integrating oncolytic viruses in combination cancer immunotherapy. Nat Rev Immunol. 2018;18(8):498–513.

7. Malvehy J, Samoylenko I, Schadendorf D, et al. Talimogene laherparepvec upregulates immune-cell populations in non-injected lesions: findings from a phase II, multicenter, open-label study in patients with stage IIIB-IVM1c melanoma. J Immunother Cancer. 2021;9(3):e001621.

8. Ribas A, Dummer R, Puzanov I, et al. Oncolytic Virotherapy Promotes Intratumoral T Cell Infiltration and Improves Anti-PD-1 Immunotherapy. Cell. 2017;170(6):1109–1119.e10.

9. Zuo S, Wei M, Xu T, et al. An engineered oncolytic vaccinia virus encoding a single-chain variable fragment against TIGIT induces effective antitumor immunity and synergizes with PD-1 or LAG-3 blockade. J Immunother Cancer. 2021;9(12):e002843.

10. Zheng M, Huang J, Tong A, Yang H. Oncolytic Viruses for Cancer Therapy: Barriers and Recent Advances. Mol Ther Oncolytics. 2019;15:234–247.

11. Macedo N, Miller DM, Haq R, Kaufman HL. Clinical landscape of oncolytic virus research in 2020. J Immunother Cancer. 2020;8(2):e001486.

12. Kawashima T, Kagawa S, Kobayashi N, et al. Telomerase-specific replication-selective virotherapy for human cancer. Clin Cancer Res. 2004 Jan 1;10(1 Pt 1):285–92.

13. Crenshaw BJ, Jones LB, Bell CR, Kumar S, Matthews QL. Perspective on Adenoviruses: Epidemiology, Pathogenicity, and Gene Therapy. Biomedicines. 2019;7(3):61.

14. Mast TC, Kierstead L, Gupta SB, et al. International epidemiology of human pre-existing adenovirus (Ad) type-5, type-6, type-26 and type-36 neutralizing antibodies: correlates of high Ad5 titers and implications for potential HIV vaccine trials. Vaccine. 2010;28(4):950–957.

15. Barouch DH, Kik SV, Weverling GJ, et al. International seroepidemiology of adenovirus serotypes 5, 26, 35, and 48 in pediatric and adult populations. Vaccine. 2011;29(32):5203–5209.

16. Huang KC, Altinoz M, Wosik K, et al. Impact of the coxsackie and adenovirus receptor (CAR) on glioma cell growth and invasion: requirement for the C-terminal domain. Int J Cancer. 2005;113(5):738–745.

17. Korn WM, Macal M, Christian C, et al. Expression of the coxsackievirus- and adenovirus receptor in gastrointestinal cancer correlates with tumor differentiation. Cancer Gene Ther. 2006;13(8):792–797.

18. Ono R, Takayama K, Sakurai F, Mizuguchi H. Efficient antitumor effects of a novel oncolytic adenovirus fully composed of species B adenovirus serotype 35. Mol Ther Oncolytics. 2021;20:399–409.

19. Vogels R, Zuijdgeest D, van Rijnsoever R, et al. Replication-deficient human adenovirus type 35 vectors for gene transfer and vaccination: efficient human cell infection and bypass of preexisting adenovirus immunity. J Virol. 2003;77(15):8263–8271.

20. Abbink P, Lemckert AA, Ewald BA, et al. Comparative seroprevalence and immunogenicity of six rare serotype recombinant adenovirus vaccine vectors from subgroups B and D. J Virol. 2007;81(9):4654–4663.

21. Su Y, Liu Y, Behrens CR, et al. Targeting CD46 for both adenocarcinoma and neuroendocrine prostate cancer. JCI Insight. 2018;3(17):e121497.

22. Elvington M, Liszewski MK, Atkinson JP. CD46 and Oncologic Interactions: Friendly Fire against Cancer. Antibodies (Basel). 2020;9(4):59.

23. Sakurai F, Nakashima K, Yamaguchi T, et al. Adenovirus serotype 35 vector-induced innate immune responses in dendritic cells derived from wild-type and human CD46-transgenic mice: Comparison with a fiber-substituted Ad vector containing fiber proteins of Ad serotype 35. J Control Release. 2010;148(2):212–218.

24. Pahl JH, Verhoeven DH, Kwappenberg KM, et al. Adenovirus type 35, but not type 5, stimulates NK cell activation via plasmacytoid dendritic cells and TLR9 signaling. Mol Immunol. 2012;51(1):91–100.

25. Johnson MJ, Petrovas C, Yamamoto T, et al. Type I IFN induced by adenovirus serotypes 28 and 35 has multiple effects on T cell immunogenicity. J Immunol. 2012;188(12):6109–6118.

26. Johnson MJ, Björkström NK, Petrovas C, et al. Type I interferon-dependent activation of NK cells by rAd28 or rAd35, but not rAd5, leads to loss of vector-insert expression. Vaccine. 2014;32(6):717–724.

27. Murakami S, Sakurai F, Kawabata K, et al. Interaction of penton base Arg-Gly-Asp motifs with integrins is crucial for adenovirus serotype 35 vector transduction in human hematopoietic cells. Gene Ther. 2007;14(21):1525–1533.

28. Ishii KJ, Coban C, Kato H, et al. A Toll-like receptor-independent antiviral response induced by double-stranded B-form DNA. Nat Immunol. 2006;7(1):40–48.

29. Blair GE, Dixon SC, Griffiths SA, Zajdel ME. Restricted replication of human adenovirus type 5 in mouse cell lines. Virus Res. 1989;14(4):339–346.

30. Jogler C, Hoffmann D, Theegarten D, Grunwald T, Uberla K, Wildner O. Replication properties of human adenovirus in vivo and in cultures of primary cells from different animal species. J Virol. 2006;80(7):3549–3558.

31. Choi YH, Lim EJ, Kim SW, Moon YW, Park KS, An HJ. IL-27 enhances IL-15/IL-18-mediated activation of human natural killer cells. J Immunother Cancer. 2019;7(1):168.

32. Umeshappa CS, Zhu Y, Bhanumathy KK, Omabe M, Chibbar R, Xiang J. Innate and adoptive immune cells contribute to natural resistance to systemic metastasis of B16 melanoma. Cancer Biother Radiopharm. 2015;30(2):72–78.

33. Nakamura T, Sato T, Endo R, et al. STING agonist loaded lipid nanoparticles overcome anti-PD-1 resistance in melanoma lung metastasis via NK cell activation. J Immunother Cancer. 2021;9(7):e002852.

34. Carreno BM, Garbow JR, Kolar GR, et al. Immunodeficient mouse strains display marked variability in growth of human melanoma lung metastases. Clin Cancer Res. 2009;15(10):3277–3286.

35. Kariya R, Matsuda K, Gotoh K, Vaeteewoottacharn K, Hattori S, Okada S. Establishment of nude mice with complete loss of lymphocytes and NK cells and application for in vivo bio-imaging. In Vivo. 2014;28(5):779–784.

36. Takeda K, Hayakawa Y, Smyth MJ, et al. Involvement of tumor necrosis factor-related apoptosis-inducing ligand in surveillance of tumor metastasis by liver natural killer cells. Nat Med. 2001;7(1):94–100.

37. Guerra N, Tan YX, Joncker NT, et al. NKG2D-deficient mice are defective in tumor surveillance in models of spontaneous malignancy. Immunity. 2008;28(4):571–580.

38. Imai K, Matsuyama S, Miyake S, Suga K, Nakachi K. Natural cytotoxic activity of peripheral-blood lymphocytes and cancer incidence: an 11-year follow-up study of a general population. Lancet. 2000;356(9244):1795–1799.

39. Gineau L, Cognet C, Kara N, et al. Partial MCM4 deficiency in patients with growth retardation, adrenal insufficiency, and natural killer cell deficiency. J Clin Invest. 2012;122(3):821–832.

40. Spinner MA, Sanchez LA, Hsu AP, et al. GATA2 deficiency: a protean disorder of hematopoiesis, lymphatics, and immunity. Blood. 2014;123(6):809–821.

41. Mandal R, Şenbabaoğlu Y, Desrichard A, et al. The head and neck cancer immune landscape and its immunotherapeutic implications. JCI Insight. 2016;1(17):e89829.

42. Shimasaki N, Jain A, Campana D. NK cells for cancer immunotherapy. Nat Rev Drug Discov. 2020;19(3):200–218.

43. Myers JA, Miller JS. Exploring the NK cell platform for cancer immunotherapy. Nat Rev Clin Oncol. 2021;18(2):85–100.

44. Marotel M, Hasim MS, Hagerman A, Ardolino M. The two-faces of NK cells in oncolytic virotherapy. Cytokine Growth Factor Rev. 2020;56:59–68.

45. El-Sherbiny YM, Holmes TD, Wetherill LF, et al. Controlled infection with a therapeutic virus defines the activation kinetics of human natural killer cells in vivo. Clin Exp Immunol. 2015;180(1):98–107.

46. Wang Y, Jin J, Li Y, et al. NK cell tumor therapy modulated by UV-inactivated oncolytic herpes simplex virus type 2 and checkpoint inhibitors. Transl Res. 2022;240:64–86.

47. Teigler JE, Kagan JC, Barouch DH. Late endosomal trafficking of alternative serotype adenovirus vaccine vectors augments antiviral innate immunity. J Virol. 2014;88(18):10354–10363.

48. Dempe S, Lavie M, Struyf S, et al. Antitumoral activity of parvovirus-mediated IL-2 and MCP-3/CCL7 delivery into human pancreatic cancer: implication of leucocyte recruitment. Cancer Immunol Immunother. 2012;61(11):2113–2123.

49. Alkayyal AA, Tai LH, Kennedy MA, et al. NK-Cell Recruitment Is Necessary for Eradication of Peritoneal Carcinomatosis with an IL12-Expressing Maraba Virus Cellular Vaccine. Cancer Immunol Res. 2017;5(3):211–221.

50. Hock K, Laengle J, Kuznetsova I, et al. Oncolytic influenza A virus expressing interleukin-15 decreases tumor growth in vivo. Surgery. 2017;161(3):735–746.

51. Xu B, Tian L, Chen J, et al. An oncolytic virus expressing a full-length antibody enhances antitumor innate immune response to glioblastoma. Nat Commun. 2021;12(1):5908.

